# Genomic characterization of *Escherichia coli* and *Enterobacter hormaechei* clinical isolates from a tertiary healthcare facility in Kenya

**DOI:** 10.64898/2026.04.13.718279

**Authors:** Sebastian Musundi, Racheal Kimani, Harrison Waweru, Patrick Wakaba, David Mbogo, Suliman Essuman, Frank Onyambu, Bernard N. Kanoi, Jesse Gitaka

**Author notes:** Current address: KEMRI Wellcome-Trust, Kilifi, Kenya (Sebastian Musundi). Current address: Nagasaki University, Japan (Patrick Wakaba). These authors contributed equally to this work (Sebastian Musundi, Racheal Kimani).

## Abstract

Extended-spectrum beta-lactamase-producing Enterobacterales such as *Escherichia coli* and *Enterobacter hormaechei* represent a growing public health challenge in clinical settings, particularly in low-and middle-income countries, due to the escalating threat of antimicrobial resistance (AMR). In this study, we aimed to identify the antibiotic resistance genes present in *E. coli* (n=4) and *E. hormaechei* (n=3) clinical isolates. Multidrug-resistant phenotypes were confirmed using disc diffusion assays against 20 antibiotics. Whole-genome sequencing of resistant isolates was performed using Oxford Nanopore Technologies. Genome assembly and analysis revealed high-risk clones, including sequence type (ST) 1193 in *E. coli* and ST78 in *E. hormaechei*. All *E. coli* isolates harbored the *bla_CTX-M_* gene in their chromosomes along with point mutations conferring resistance to fluoroquinolones, while *E. hormaechei* isolates encoded *bla_ACT_* in their chromosomes. Additionally, both species carried plasmids with multiple antibiotic resistance genes, including *bla_OXA_* and *bla_TEM_*, co-located with metal resistance operons, indicating the potential for horizontal gene transfer. BLAST analysis revealed high sequence similarity between the plasmids identified in clinical isolates and those previously recovered from environmental sources, highlighting the role of environmental reservoirs in AMR dissemination. Notably, no carbapenem resistance genes were detected in any isolate. These findings underscore the growing threat posed by multidrug-resistant Enterobacterales in clinical settings and emphasize the urgent need for strengthened infection prevention and control measures to mitigate AMR spread.

## Introduction

Over the past 70 years, treating and controlling bacterial diseases has greatly relied on antibiotics [1]. However, the rise in antimicrobial resistance (AMR) in the past decades has become a significant global public health challenge [2]. In 2019 alone, the World Health Organization estimated 1.27 million deaths from antibiotic-resistant bacteria [3]. Although all regions across the world and all income levels are affected by antimicrobial resistance, the impact is exacerbated in low-and-middle-income countries where there is the indiscriminate use of antibiotics, poor surveillance, and a lack of stewardship programs [4,5].

Among the most concerning pathogens identified by the World Health Organization as critical for AMR management are extended-spectrum beta-lactamase-producing *Enterobacterales* (ESBL-E). These pathogens are resistant to third-generation cephalosporins and last-line antibiotics like carbapenems[6]. In Kenya, there is increasing evidence of resistance to cephalosporins[7,8], a worrying trend given that beta-lactams are essential for treating most ESBL-E*-*associated infections. Although carbapenems can be used to treat ESBL-E, their use in Kenyan healthcare facilities is limited due to their high cost and infrequent prescription [9]. Moreover, the potential for carbapenem-resistant *Enterobacterales* to emerge is a growing concern, exacerbated by the limited availability of alternative treatments like ceftazidime-avibactam.

Resistance in ESBL-E arises through various mechanisms, including the production of beta-lactamase enzymes that cleave the beta-lactam ring, efflux pumps, and alteration in the penicillin-binding sites [10]. In response to beta-lactam exposure, AmpC resistance can also be induced on the bacterial chromosomes [11]. Mobile genetic elements such as plasmids, integrons, and transposons facilitate the spread of ESBLs and other antibiotic-resistance genes (ARGs) horizontally. Among the most prevalent ESBL genes against cephalosporins is bla_CTX-M_ [10].

Routine genomic surveillance offers a valuable opportunity to investigate the spread of ARGs among ESBL-E providing essential data to inform infection and prevention control measures. Next-generation sequencing technologies have greatly facilitated this approach, with short-read sequencing currently being the gold standard for whole genome sequencing due to its lower error rate and high accuracy [11]. However, short-read technologies face notable challenges in assembling complete bacterial genomes, particularly in resolving repetitive regions and reliably reconstructing plasmids [12]. Since 2014, Oxford Nanopore Technologies (ONT) has developed and released long-read sequencing platforms capable of generating reads that span these repetitive regions, providing an approach that could generate complete bacterial genomes and resolve mobile genetic elements [13,14]. Moreover, ONT offers portable devices with short turnaround times and low initial investment, which aids in rapid, cost-effective genomic surveillance and is suitable for resource-limited settings. The availability of full genomes greatly improves our understanding of bacterial structure and function, evolution, and identification of mobile genetic elements and ARGs.

Within healthcare settings, *Enterobacterales Escherichia coli* and *Enterobacter hormaechei* have become increasingly important pathogens due to their production of ESBLs, complicating the treatment of infections with cephalosporins and other antibiotics. *E. coli* is the leading cause of urinary tract infections[15], while *E. hormaechei* is an important nosocomial and opportunistic pathogen associated with urinary tract infections, colitis, pneumonia, and other diseases [16]. In this study, we report on ESBL-E *E. coli* and *E. hormaechei* isolates from clinical isolates sequenced using ONT. Herein, we show the presence of bla_CTX_ in *E. coli* isolates and bla_ACT_ encoded in *E. hormaechei* chromosome. We also highlight the detection of high-risk clones ST1193 in *E. coli* and ST78 in *E. hormaechei* and the presence of multiple plasmids containing multiple ESBLs genes and metal resistance operons. These plasmids facilitate the horizontal transfer of ARGs, contributing to the spread of resistance and further complicating treatment options within healthcare settings.

## Materials and Methods

### Ethics statement

This study was approved by the Institutional Scientific and Ethics Research Committee (ISERC) of Mount Kenya University (MKU/ERC/1687) and licensed by the National Commission for Science, Technology, and Innovation (NACOSTI) (NACOSTI /P/21/7678). Before enrolment, written informed consent was obtained from each participant or their legal guardian.

### Bacterial isolates and DNA extraction

The bacterial isolates (n=202) used in this study were collected between March and November 2021 from patients at Thika Level V Hospital. The isolates were previously cultured and stored in frozen glycerol stocks at the Mount Kenya University research laboratory. The isolates had been previously identified by VITEK (Biomerieux). Antimicrobial susceptibility testing was carried out using the Kirby-Bauer disk diffusion method. Isolates were tested against the following panel of antibiotics: carbapenems (imipenem and meropenem) (10µg), cephalosporins (cefuroxime, ceftriaxone, cefepime, and ceftazidime, cefotaxime) (30 µg), cephamycins (cefoxitin(30 µg), monoamides (aztreonam (30 µg), quinolones (ciprofloxacin and levofloxacin) (5 µg), aminoglycosides (gentamicin(10 µg) and amikacin(30 µg)), penicillins (piperacillin (30 µg), and ampicillin (10 µg), tetracyclines (30 µg (Minocycline (30 µg) and tetracycline (30 µg)) and sulfonamides (sulfamethoxazole combined with trimethoprim (25µg)). Combinations of drugs used were amoxicillin-clavulanate (30 µg) and piperacillin-tazobactam (110 µg) (Oxoid, England). Diffusion diameters were interpreted based on the Clinical Laboratory Standards and Institute (CLSI 2020) guidelines [17] as resistant (R), susceptible (S), or intermediate (I). *E. coli* ATCC 25922^®^ and *Klebsiella pneumoniae* ATCC 700603^®^ were used as a reference for the interpretation of the data based on the guidelines of CLSI.

*Escherichia coli* (n=4) and *Enterobacter hormaechei* (n=3) isolates that exhibited multidrug resistance in the disc diffusion assay were cultured on nutrient agar for 18 hours at 37^0^C. Following incubation, individual colonies were selected and suspended in 1X PBS to create a bacterial suspension. Genomic DNA was then extracted using Zymogen® bacterial and fungal DNA extraction kit. The quality and quantity of genomic DNA was then assessed using the NanoDrop and Qubit 3.0 fluorometer.

### ONT library preparation and sequencing

DNA library preparation was carried out using the SQK-LSK109 ligation sequencing kit. The fragmented DNA was first repaired using the NEBNext FFPE DNA Repair Mix and NEBNext Ultra II End Repair/dA-Tailing Module (New England BioLabs). Subsequently, individual barcodes were incorporated into the dA-tailed DNA using the EXP-NBD104 and EXP-NBD114 native barcoding expansion kit following the ONT protocol with NEB Blunt//TA Ligase Master Mix (New Englands Biolabs). Barcoded DNA samples were then equimolarly pooled, and adapters were attached using the Quick T4 DNA Ligase Quick Ligation Module (New Englands Biolabs). Before sequencing, the number of active pores on the flow cell R9.5(FLO-MIN106) was assessed, and equimolar pooling of samples was performed. Finally, sequencing was then carried out on the MinKNOW for 48 hours.

### Quality control

Basecalling of the raw Fast5 files produced after sequencing was performed using Guppy version 6.4.6 with the high accuracy mode option without quality filtering options. The multiple FASTQ files were merged into one and demultiplexed using the Guppy barcoder [18]. FastQC was used to check the quality of the sequences. Afterward, Porechop (https://github.com/rrwick/Porechop) was used to trim adapters, while Filtlong (https://github.com/rrwick/Filtlong) was used to filter reads with less than 500 base pairs. These steps contribute to the generation of high-quality sequences for downstream analysis.

### De novo assembly, polishing, and annotation

Flye. 2.9.1 (https://github.com/fenderglass/Flye) generated the draft assembly from the processed FASTQ reads [19]. The resulting draft assembly was then indexed and mapped against the individual reads using BWA-MEM (https://github.com/bwa-mem2/bwa-mem2) [20]. The generated SAM files were sorted, indexed, and converted to BAM format using Samtools (https://github.com/samtools/samtools). A three-step polishing approach was employed to improve the assembly quality. Four rounds of Racon (https://github.com/isovic/racon) were first utilized to polish the assembly using the mapped nanopore reads [21]. Subsequently, Medaka 1.7.2 (https://github.com/nanoporetech/medaka) and Homopolish were used to improve polish reads. The quality of the draft genome assembly was assessed using Quast (https://github.com/ablab/quast), and the completeness of the draft assembly was evaluated using BUSCO (https://github.com/WenchaoLin/BUSCO-Mod). Finally, the draft genome was annotated using Prokka (https://github.com/tseemann/prokka), which predicts coding sequences and other genomic features from bacterial genomes [22].

### Species identification

PubMLST was used to identify the specific bacterial species from the assembled contigs [23]. In addition, we also used the average nucleotide identity (ANI) score to validate our species classification results from PubMLST. ANI compares the nucleotide identity in all orthologous genes shared within the same species [24]. Organisms belonging to the same species have an ANI>95%. We used *Escherichia coli* str. K-12 substr. MG1655 (NC_000913.3) and *Enterobacter hormaechei* (CDC Enteric Group 75) (CP077308.1) as references to determine the ANI scores for *E. coli* and *E. hormaechei* isolates, respectively. For *E. coli*, we used Clermon typing to classify the isolates into individual phylogroups [25] and SerotypeFinder 2.0 to identify individual serovars [26].

### In-silico prediction of ARGs and plasmids

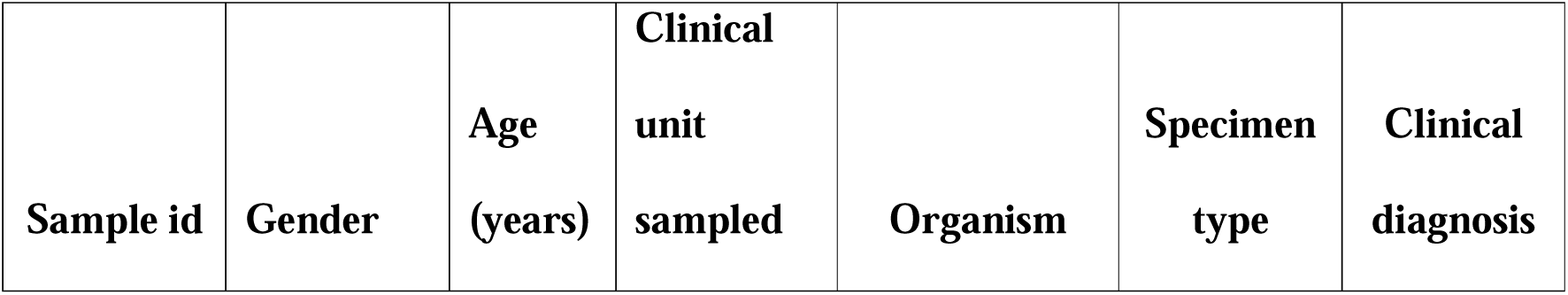

Prediction of antimicrobial resistance genes was conducted using Resfinder version 4.6.0, which had a minimum length of 60% and an identity threshold of 90% on the assembled bacterial chromosome. PlasmidFinder was used to identify plasmid replicons in contigs [27]. Additionally, we used the mob-recon tool present in the MOB-suite to reconstruct contigs from the assembly [28]. The reconstructed plasmid contigs were further subjected to a nucleotide BLAST against the core nucleotide database, and similar plasmids were visualized using Genofig [29]. Proksee was also utilized to visualize and characterize plasmid sequence [30]

## Results

### Patient characteristics

The demographic and clinical information from where the seven isolates were isolated is summarized in Table 1. Most isolates (5/7) were from the renal unit of patients diagnosed with end-stage renal disease (ESRD), primarily adults aged 38-63 years. The remaining two isolates were from a 21-year-old male following a trauma incident (surgical ward) and a 91-year-old female diagnosed with congestive heart failure (CHF) (intensive care unit). Among the isolates, 4 were E. *coli* (urine and stool), and 3 were *E. hormaechei* (pus and urine).

**Table 1.** Summary of patient demographics and clinical diagnoses for isolates used in the study. Samples were from the surgical ward, intensive care unit (ICU), and renal unit, while the clinical diagnosis included trauma from a root traffic accident (RTA), congestive heart failure, and end-stage renal disease (ESRD).

|  |  |  |  |  |  |  |
| --- | --- | --- | --- | --- | --- | --- |
| S03 | Male | 21 | Surgical ward | *E. hormaechei* | pus | Trauma/<br>RTA |
| S08 | Female | 91 | ICU | *E. hormaechei* | urine | CHF |
| S09 | Male | 63 | Renal Unit | *E. hormaechei* | urine | ESRD |
| S24 | Male | 63 | Renal Unit | *E. coli* | urine | ESRD |
| S26 | Female | 38 | Renal Unit | *E. coli* | urine | ESRD |
| S27 | Female | 38 | Renal Unit | *E. coli* | urine | ESRD |
| S29 | Female | 57 | Renal Unit | *E. coli* | stool | ESRD |

### Antimicrobial susceptibility tests reveal multidrug resistance in *E. coli* and *E. hormaechei* isolates

All *E. coli* isolates consistently resisted Ceftriaxone, Cefotaxime, and Ampicillin, while Ciprofloxacin and Tetracycline showed resistance, particularly in urine samples. As both susceptible and resistant profiles were observed, mixed resistance profiles were observed for Sulfamethoxazole/Trimethoprim and levofloxacin. For the *E. hormaechei* isolates, resistance was observed to cephalosporins Cefotaxime, Ceftriaxone, and fluoroquinolones Ciprofloxacin and levofloxacin while susceptibility was observed for Sulfamethoxazole/Trimethoprim (SXT) and Levofloxacin in some isolates. All *E. coli* and *E. hormaechei* isolates were susceptible to the carbapenems, Imipenem, and Meropenem. The results are summarized in S1 Table.

### Multidrug resistance *E. coli* isolates with chromosomal mutations and composite transposons mediating antibiotic resistance

We assembled four *E. coli* genomes with the length of the largest contig ranging from 1,042,887 to 4,948,395 bp. The average nucleotide identity (ANI) for all isolates was >95%, and all isolates fell under phylogroup B2 based on Clermon typing (Table 2). MLST analysis classified two isolates as ST1193, whereas the remainder were ST53 and ST12270 (Table 2). The most common serovar was O75:H5, present in three of four isolates, while the remaining serovar was O25:H4 (Table 2). We also detected the presence of multiple antimicrobial resistance genes (ARGs) in the chromosomal region of the *E. coli* isolates within the insertion sequences. The extended-spectrum beta-lactamase (ESBL), *bla_CTX-M-15,_* was present in the chromosomes of all isolates. Additionally, three of the four isolates (S24, S26 and S27) had the ESBL *bla_OXA-1_*, and aminoglycoside resistant genes *aac(6’)-Ib-cr* and *aac(3)-Iia* (Table 2). We further identified chromosomal point mutations that mediate AMR genes in all *E. coli* isolates. These mutations were primarily found in the genes encoding DNA gyrase A (*gyrA*), topoisomerase IV *parC,* and *parE* and β-subunit of RNA polymerase *rpoB* (Table 2). Point mutations in *gryA, parC,* and *parE* are associated with resistance to fluoroquinolones, while mutations *rpoB* confers resistance to rifampicin.

**Table 2.** Genomic characteristics of the assembled *E. coli* isolates. The table shows the individual sequence types based on MLST classification, phylogroups, average nucleotide identity against E. coli reference, the length of the largest contigs, serovar classification, and resistance genes located within the chromosomal regions. Chromosomal mutations mediating antibiotic resistance are illustrated as point mutations in specific genes.

| Sample ID | Sequence type | Phylogroup | Average nucleotide identity | Largest contig length | Serovar | Resistance genes (composite transposons) | Chromosomal point mutations mediating AMR |
| --- | --- | --- | --- | --- | --- | --- | --- |
| S24 | ST12270 | B2 | 96.366 | 3,166,760 | O25:H4 | *bla<sub>CTX-M-15</sub>*, *bla<sub>OXA-1</sub>*,<br>*aac(6')-Ib-cr* | *parC* (p.S80I), *parC*(p.E84V) |
| S26 | ST53 | B2 | 96.481 | 1,042,887 | O75:H5 | *bla<sub>CTX-M-15</sub>*, *aac(3)-Iia*, *catB3*,<br>*bla<sub>OXA-1</sub>*, *aac(6')-Ib-cr* | *parC* (p.S80I), *rpoB*(p.P564L),<br>*gyrA*(p.S83L), *gryA*(p.D87N),<br>*parE*(p.L416F) |
| S27 | ST1193 | B2 | 96.87 | 4,948,395 | O75:H5 | *aac(3)-Iia*, *aac(6')-Ib-cr*,<br>*bla<sub>CTX-M-15</sub>*, *bla<sub>OXA-1</sub>*, *catB3* | *gyrA*(p.S83L), *gryA*(p.D87N),<br>*parE*(p.L416F) |
| S29 | ST1193 | B2 | 96.52 | 1,854,372 | O75:H5 | *aac(3)-Iia*, *bla<sub>CTX-M-15</sub>* | *gyrA*(p.S83L), *gryA*(p.D87N) |

### Plasmids isolated from *E. coli* contained multiple ARGs and metal resistance operons

We isolated multiple plasmids in three out of the four *E. coli* isolates. The plasmid in isolate S26 was 92,194bp long and classified under the IncFIB replicon. Blast analysis of this plasmid, subsequently labeled pS26, against the core nucleotide database revealed multiple similar plasmids with a query cover >=95% and percentage identity >=95% (Fig 1A). Phylogenetic analysis of these plasmids revealed a monophyletic group comprising plasmids sampled in China in 2019 and Switzerland in 2021 (Fig 1B). The separate monophyletic clusters suggest potential local dissemination of the plasmid in Switzerland. It may indicate interactions between clinical and environmental reservoirs in China as the samples were isolated from stool and sink drain. The plasmid isolated from Kenya exhibited genetic similarity to other plasmids from China and Switzerland compared to Australia. Plasmids isolated from China and Switzerland likely served as the potential source of global dissemination.

**Fig 1.**
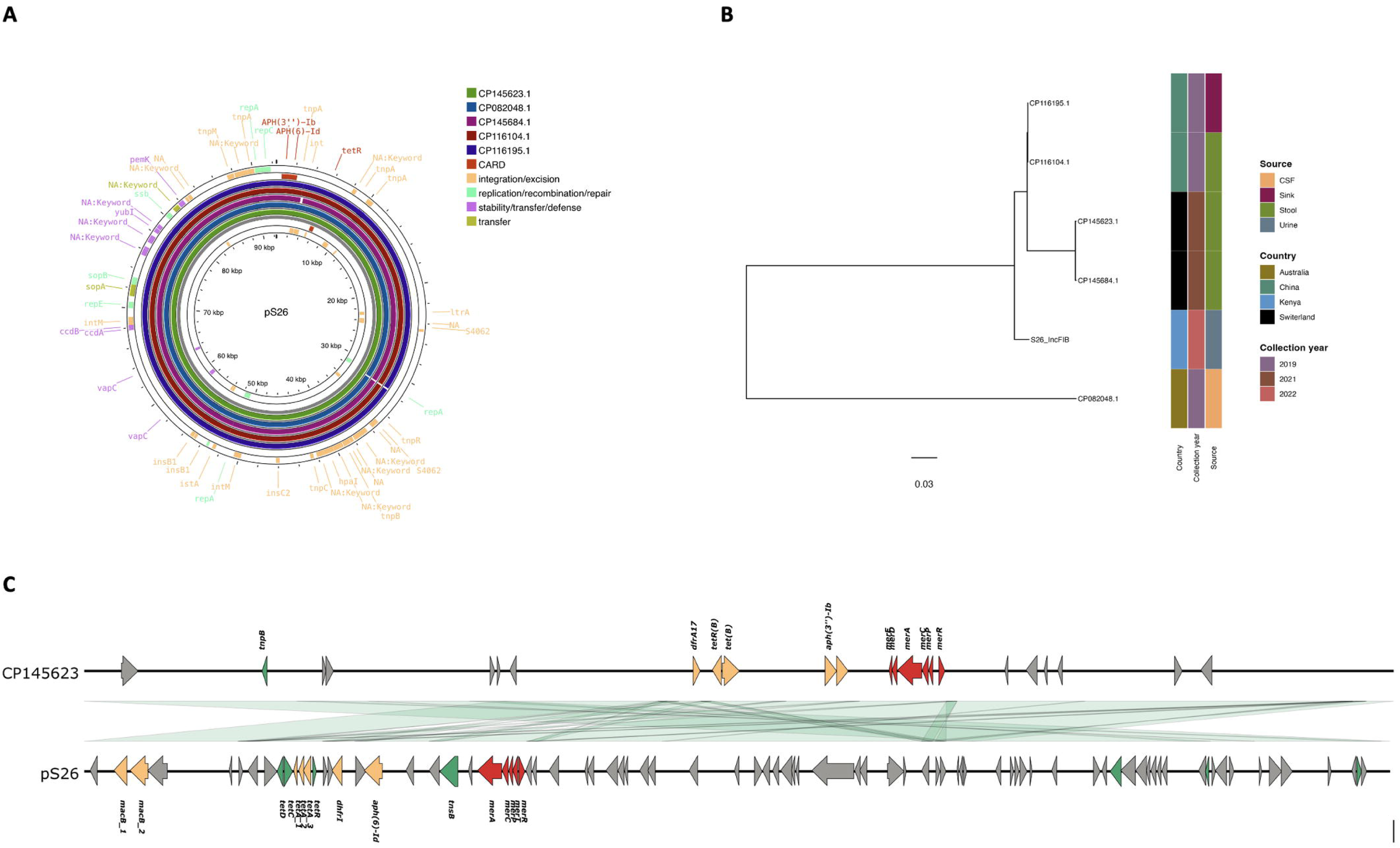
Genomic analysis of plasmid sequences retrieved from an *E. coli* isolate. A) Comparison of *E. coli* plasmid with others retrieved from NCBI. The grey circle indicates plasmids pS26, while the four circular rings afterward represent sequence similarity with other plasmids retrieved from NCBI. The red color indicates the presence of ARGs retrieved from the CARD database. MobileOG-db was used to identify genes mediating integration or excision, recombination, defense, or transfer of mobile genetic elements. B) Maximum likelihood tree showing the relationship between pS26, replicon lncFIB, and plasmids with high similarity (>95%) and coverage (>95%) from NCBI. The samples were annotated based on their country of origin, collection year, and sample source/site. C) pS26 similarity with plasmid CP145623.1 highlights ARGs in orange, metal resistance genes in red, hypothetical protein in grey, and insertion sequences and transposons in green. The green color between the two plasmids shows a color gradient with the minimum sequence similarity between the identified genes at 90%.

Furthermore, detecting the plasmid across diverse sample sites, including urine, stool, sink drain, and cerebrospinal fluid, underscores its potential for widespread dissemination and persistence in multiple ecological niches. Further analysis of this plasmid revealed the presence of ARGs mediating resistance against macrolides (*MacB*), aminoglycosides (*aph(6)-Id)*, tetracycline (*tetA, tetC, tetD, tetR*), and trimethoprim (*dhfr1*) within a transposon region. In addition, the mercuric resistance operon genes (*mer A, merR, merT, merP*, and *merC*) responsible for the transport and detoxification of mercury were also present in plasmid CP145623.1 (Fig 1C)

A plasmid isolated from S27 and labeled pS27 was 92,199bp long and showed high similarity with plasmid CP082825.1 isolated from urine in Kazakhstan in 2021. Similar to the previous plasmid, multiple ARGs against tetracycline (*tetA, tetC, tetD,tetR*), aminoglycoside (*aph(6)-Id*), and macrolides (*MacB*) were identified in the plasmid as well as the presence of mercuric resistance operon (*mer A, merR, merT, merP,* and *merC*) (Fig 2A). We also identified four plasmids in S29 of lengths 56,596bp, 92401bp, 55,769bp, and 287,498bp, which were named pS29_001, pS29, pS29_002, and pS29_003, respectively. The plasmid pS29_001 was similar (>95%) to plasmid CP120877.1 isolated from China in urine in 2019. Moreover, it contained multiple ARGs conferring resistance to beta-lactams(*AmpC, bla_OXA-1_*), aminoglycosides *(aacA4*), and chloramphenicol (*catB*). This plasmid also possessed a multidrug transporter efflux pump (*ermE*) and phage shock protein operon (*pspF, pspA, pspB, pspC,* and *pspD*) (Fig 2B). pS29 was similar to a stool sample isolated in Switzerland in 2021 CP145623 with a percentage similarity of 99.88%. Also, it contained multiple resistance genes against aminoglycosides (*aph(6)-Id*), tetracycline (*tetA, tetC, tetD, tetR*), and macrolides (*MacB*) as well as having a mercuric operon (*mer A, merR, merT, merP,* and *merC)*. The third plasmid, pS29_002, was 99.42% identical to plasmid AP027596.1 isolated from feces in Ecuador in 2019 but showed less coverage (50%), while the fourth plasmid, pS29_003 showed 99% percent similarity and 80% coverage with CP045203.1, from a rectal sample from Israel in 2010. Analysis of the ARGs in these two plasmid samples using the Resfinder database showed resistance to aminoglycosides (*aph(6)-Id, aadA1)*, beta-lactams (*bla_TEM-1B_, bla_DHA-1_, bla_TEM-148_* and *bla_OXA-1_*), sulfonamides (*sul1*), macrolides (*mph(A))*, chloramphenicol (*catB3*), quinolones (*qnrB4*) and trimethoprim (*dfrA1*)(Supplementary Table 2). Based on the above analysis, two plasmids carried multidrug resistance genes and metal operon resistance genes, while another plasmid contained phage shock protein operon in combination with multiple ARG

**Fig 2.**
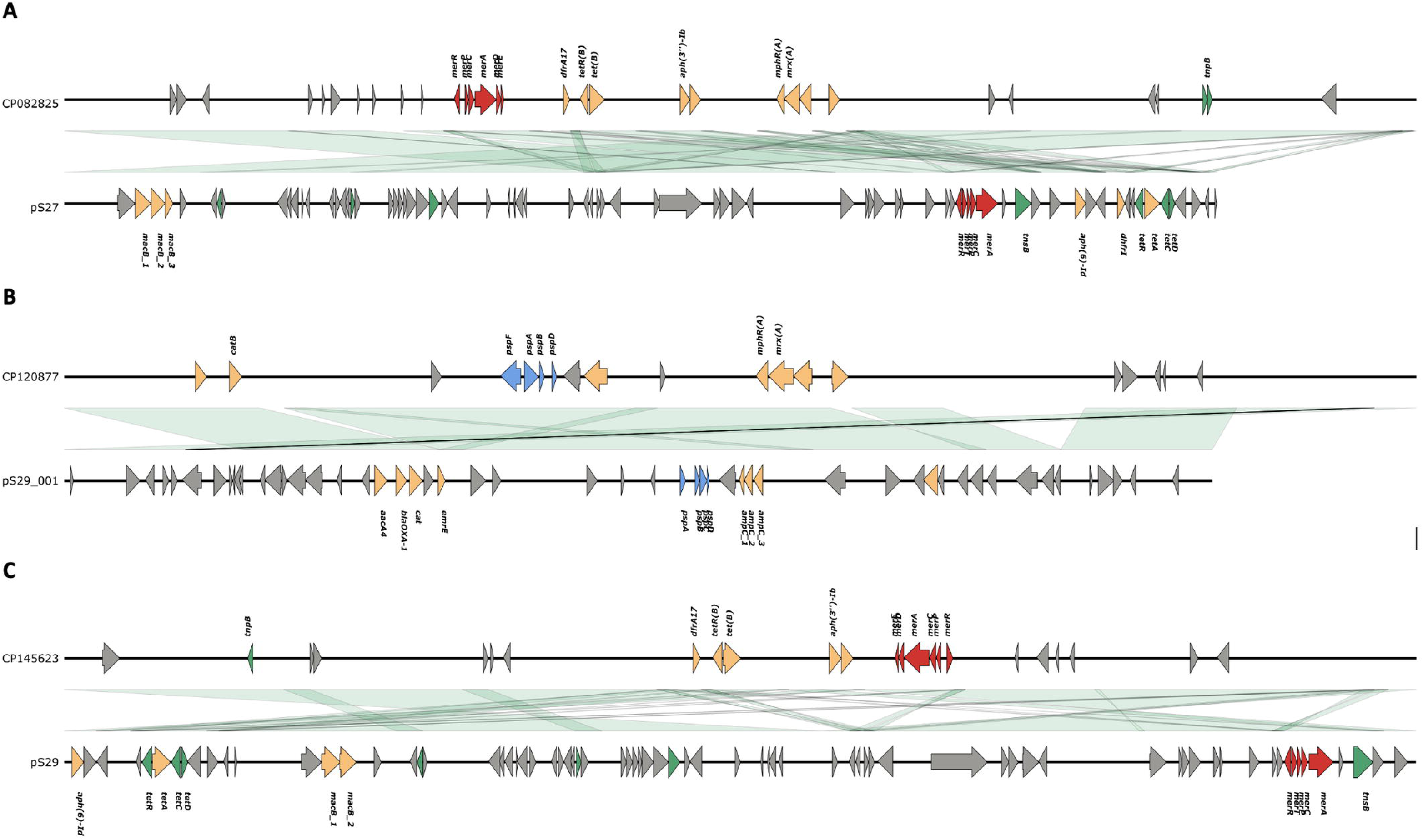
Plasmid sequences isolated from *E. coli* with high similarity to previously deposited plasmids in NCBI. A) pS27 was identical to CP082825 isolated from Kazakhstan, B) pS29_001 similar to CP120877.1 isolated from China, and C) pS29 identical to CP145623 isolated from Switzerland. The individual plasmids were annotated using Prokka. The red color indicates metal resistance operons, the grey color suggests hypothetical proteins, the green represents insertion sequences and transposons, and the blue color indicates phage-related proteins. The green color between the two plasmids shows a color gradient with the minimum sequence similarity between the identified genes at 90%.

### *Enterobacter hormaechei* chromosome encodes bla_ACT_ and plasmids carrying multiple ARGs and metal resistance operons

We further assembled three *Enterobacter hormaechei* isolates, S03, S08, and S09, and obtained chromosomal genome sizes of 4,807,223bp, 4,668,797bp, and 4,579,703bp, respectively. The three isolates had an average nucleotide identity above 94% compared to the reference, confirming their identity as *Enterobacter hormaechei*. All three isolates possessed the AmpC beta-lactamase genes, *bla*_ACT,_ while S08 and S09 had the presence of the fosA, which confers resistance to fosfomycin encoded in their chromosome. MLST analysis showed that isolate number S03 belonged to ST88 while isolate S08 belonged to ST98. Isolate number S09 belonged to the high-risk clone ST78.

Isolate designated as S03 contained a plasmid of 216,294 bp, 99.97% identical to a plasmid isolated in the USA in 2013 from the feedlot (CP104113). The plasmid labeled pS03 contained multiple ARGs located within transposons, including those conferring resistance to beta-lactamase (bla_OXA-1_, *ampC*), chloramphenicol (*catB*), tetracycline (*tet, tetR*), and aminoglycosides (*aph(6)-ld, aacA4*) (Fig 3A). An additional plasmid in isolate number S03 of approximately 109,591 bp was 99.42% identical to a plasmid CP143700.1 isolated in Taiwan in 2024. Analysis of this contig on the Resfinder database revealed the presence of the bla_CTX-M-15_ (S2 Table). Isolate number S08 had plasmid pS08, which was 80,264 bp and 99.98% identical to AP021911.1, isolated from a wastewater treatment plant in Japan in 2017. The primer pS08 contained genes conferring resistance to beta-lactamase (*bla_TEM,_bla_CTX-M-1_*), aminoglycosides (*aph(6)-ld)*, and diaminopyrimidines (dhfr1), as well as a mercuric operon (mer A, merR, merT, merP, and *merC*) (Fig 3B). Lastly, the high-risk clone ST78 contained a 96,831bp plasmid labeled pS09, which was 99.87% identical to CP095178.1 isolated in China in 2019. ARGs present contained resistance genes against beta-lactams (*bla_0XA-1_, bla_CTX-M-1_, bla_TEM-12_*), aminoglycoside (*aacA4, aph(6)-ld*), tetracycline (*tetR, tetA*), diaminopyrimidine (*dhfr1*) and chloramphenicol (*catB*) (Fig 3C). Unlike the previous plasmid pS08, which showed the mercuric operon, pS09 contained the arsenic operon (Fig 3C). Plasmids isolated from *E. hormaechei* showed the presence of multiple ARGs within the same plasmid and metal resistance operon.

**Fig 3.**
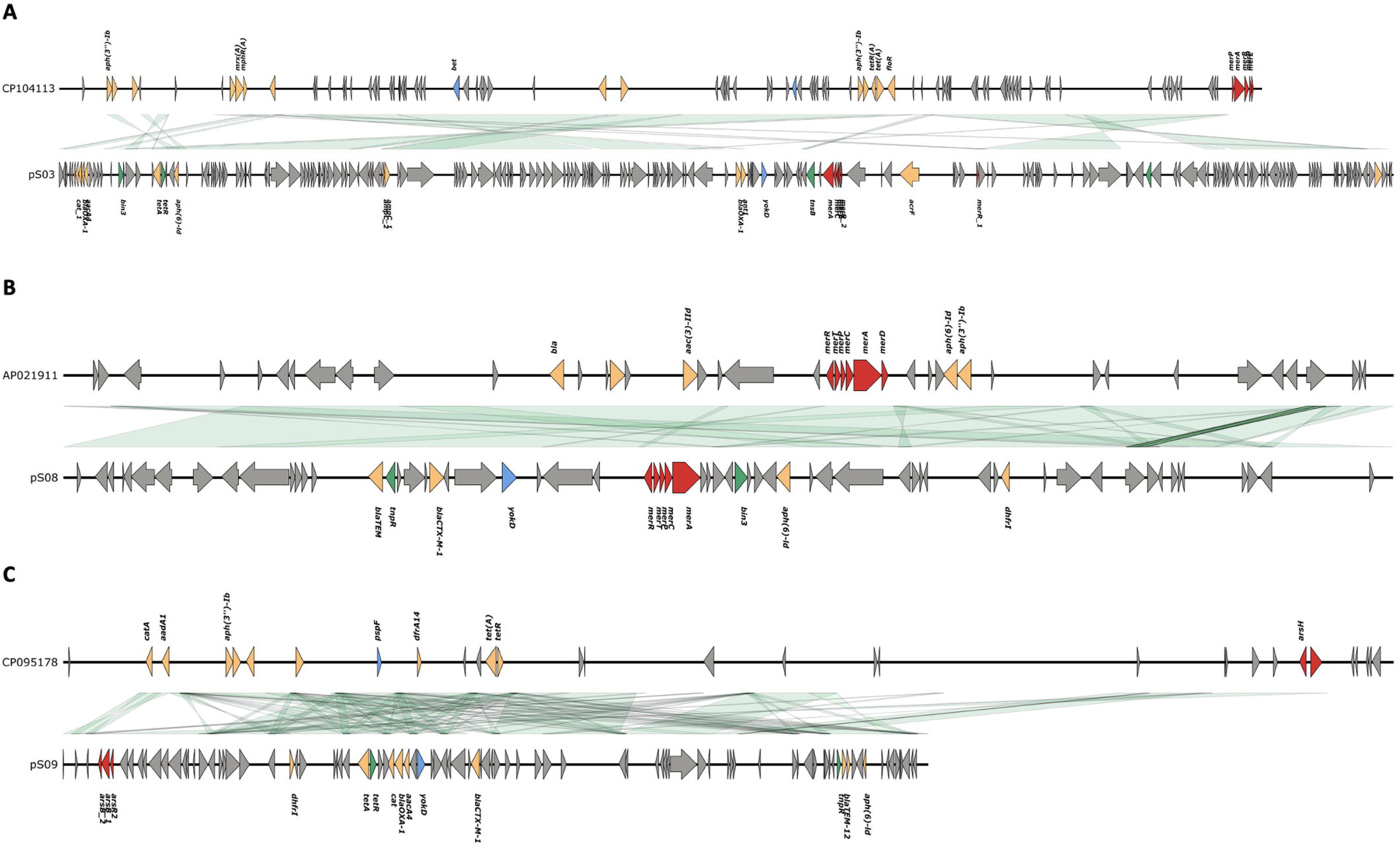
Comparison of plasmids isolated from *E. hormaechei* isolates with those in NCBI. A) pS03 similar to plasmid CP104113 isolated from a feedlot in the USA, B) pS08 identical to AP021911 isolated from a wastewater plant in Japan, and C) pS09 similar to CP095178 isolated from China. The grey color across all plasmids indicates hypothetical genes after annotation using Prokka; red indicates metal resistance operon, orange indicates ARGs, green represents insertions or transposons, and blue indicates phage sequences.

## Discussion

In clinical settings, ESBL-producing Enterobacterales, particularly *E. coli,* and *E.hormaechei*, are major contributors to hospital-acquired infections (HAI) [31]. Their isolation in clinical samples underscores their role as nosocomial pathogens. Trauma patients, particularly those with open wounds or undergoing catheterization, face a higher risk of acquiring infections from resistant pathogens such as *E. hormaechei*. Similarly, patients with ESRD are particularly vulnerable to HAI due to multiple factors, including the use of invasive devices such as dialysis catheters, frequent hospitalization for dialysis [32], and weakened immune systems, which make them prone to opportunistic infections from *E. coli* and *E. hormaechei*. These factors make ESRD patients more susceptible to infections caused by *E. coli* and *E. hormaechei,* which are often resistant to first-line antibiotics and present a challenge to effective treatment.

To better understand the resistance mechanisms driving these HAIs, we isolated and sequenced *E. coli* and *E.hormaechei* from clinical samples using nanopore sequencing. Two of the four *E.coli* isolates from this study were classified as ST1193, a global multidrug high-risk clone implicated as an essential cause of bloodstream and urinary tract infections. This high-risk clone is particularly concerning in the context of ESRD patients due to its ability to colonize and persist in hosts for long periods and cause recurrent infections [33]. Similar to other ST1193 isolates, our two isolates showed the presence of chromosomal point mutations in *gyrA* at S83L and D87N against fluoroquinolones, complicating standard antibiotic treatment. Furthermore, it belonged to the virulent phylogroup B2 with the O-type 75[34] and integrated *bla_CTX-M-15,_ bla_OXA-1_*, and other ARGs in the chromosome of some of the ST types, including ST1193. This increases its heritability and stability, making resistance more persistent in bacterial populations and contributing to the long-term dissemination of the resistant trait in clinical settings. Integrating genes conferring resistance of beta-lactams, especially bla_CTX-M_ and other ARGs mediated by insertion sequences carried by transposition units [30], will likely result in a multidrug resistance island that complicates available treatment options.

Following the analysis of *E. coli* isolates, we also isolated ST78 *E*. *hormaechei*, a primary driver for the transmission of carbapenemase-producing *E. hormaechei* (CPEH), from one of the isolates. ST78 is currently recognized as a high-risk clone because it can acquire multiple plasmids carrying ARGs. This strain is particularly concerning in the context of HAI, where resistant pathogens are more likely to affect vulnerable patients, such as those with ESRD, as in the current study. Notably, one of the *E. hormaechei* isolates came from a 91-year-old ICU patient and another from a 21-year-old trauma patient in the surgical ward, both carrying the bla_ACT_ gene, which confers resistance to penicillins and first-and second-generation cephalosporins [35,36]. The presence of bla_ACT_ in these distinct patient populations is concerning as it highlights the prevalence of resistance across different hospital units. Although each isolate also harbored a plasmid carrying a gene responsible for beta-lactam resistance, plasmids isolated from ST78 carried three beta-lactam conferring genes -*bla_OXA-1_*, *bla_TEM_, and bla_CTX-M_,* but lacked any resistance genes to last-line antibiotics such as carbapenem. Nevertheless, multiple ARGs on these plasmids highlight the potential for horizontal gene transfer and contribution to the spread of multidrug-resistant strains, further complicating treatment strategies.

Mobile genetic elements, particularly plasmids identified in both *E. coli* and *E. hormaechei* isolates, carried the most significant number of ARGs genes conferring resistance to multiple antibiotics, including tetracyclines, macrolides, chloramphenicol, diaminopyrimidines as well as beta-lactams. Plasmids are crucial in disseminating AMR through horizontal gene transfer [37], enabling rapid spread of resistance across bacterial populations. ST1193 isolates in *E. coli* contained additional ESBL, including *bla_TEM_, bla_DHA,_* and *bla_OXA,_* alongside other ARGs, further emphasizing the pivotal role of plasmids in contributing to multidrug resistance. Interestingly, some plasmids isolated from clinical samples were identical to plasmids previously identified in environmental sources, underscoring the environment’s significant role in the spread and transmission of multidrug resistance strains via conjugation, transduction, or transformation [38]. The presence of these plasmids in clinical and environmental settings highlights the need for infection and prevention control measures that target both settings. Furthermore, *E. coli* and *E. hormaechei* plasmids also carried metal resistance operons, particularly those conferring resistance to mercury and arsenic. The presence of these operons promotes mercury and arsenic tolerance, allowing for co-resistance when heavy metal and antibiotic resistance genes are located on the same mobile genetic element [39]. The co-selective pressure of these heavy metals remains a significant contributor to the dissemination and persistence of ARGs in environmental settings.

The presence of identified high-risk clones ST1193 and ST78 in *E. coli* and *E. hormaechei* in clinical isolates, as well as the presence of ESBLs and multiple ARGs, could provide insights on empirical therapy decisions, infection control, and antibiotic stewardship programs to be implemented in healthcare settings. Early identification of ARGs, especially bla_CTX-M_ and bla_OXA,_ and the associated plasmids allows clinicians to refine empirical antibiotic choices, minimizing the risk of ineffective treatments and promoting narrow-spectrum agents where possible. Such measures align with good antibiotic stewardship programs that emphasize the need to delay antibiotic initiation until the availability of diagnostic test results [40]. The alternative use of broad-spectrum antibiotic treatment without testing high-risk clones will likely increase selection pressure, resulting in the selection of MDR strains. Applying infection control and preventive measures would also likely reduce incidences of community infections and nosocomial infections. Already, several initiatives in LMICs, such as the establishment of a national AMR surveillance system across multiple sentinel hospital sites to monitor AMR spread [41], the development of regulatory frameworks for the prescription of antibiotics [42], and implementation of antibiotic stewardship programs [43] are some of the effective measures in place to reduce the threat posed by AMR.

While our study highlights the presence of ESBL-producing Enterobacterales and the associated ARGs, it is limited by several factors. The relatively small number of isolates collected from a single tertiary facility restricts the generalizability of our findings. Additionally, the absence of genomic data from the surrounding environmental isolates constrains any broader contextual transmission analysis between the hospital units. Given the identification of high-risk clones linked to multidrug resistance, future large-scale multicenter genomic surveillance studies are needed to understand the spread and evolution of ARGs. Meanwhile, it’s essential that the routine phenotypic confirmation approaches, such as the double disc synergy test, carbapenemase Nordmann–Poirel Test, and modified carbapenem inactivation method, continue to be applied within clinical settings to strengthen the genotype-phenotype link and guide targeted interventions.

## Conclusion

ESBL-E pose a public health threat to clinical settings in low and middle-income countries such as Kenya. Our findings indicate the presence of multidrug-resistant strains of *E. coli* and *E. hormaechei* isolated from clinical settings against commonly used cephalosporins.

Genomic analysis also reveals the presence of high-risk multidrug clones ST1193 and ST78 in *E. coli* and *E. hormaechei,* respectively, and plasmids facilitating the horizontal transfer of ARGs. Although we did not observe carbapenem-resistant genes, our findings underscore the critical need for genomic surveillance to monitor and address the emerging threat posed by ESBL-producing pathogens. To combat AMR, we strongly recommend implementing antimicrobial stewardship programs alongside infection prevention and control measures in healthcare settings. These measures are essential in curbing the spread of resistant infections and ensuring the effectiveness of available antibiotics.

## Acknowledgments

We appreciate the technical assistance of the study volunteers and research teams from Thika Level V referral hospital in Kiambu County and the research team at the Center for Malaria Elimination at Mount Kenya University.

## Supporting information

**S1 Table. Antimicrobial susceptibility results for the *E. coli* and *E. hormaechei* isolates.** Test carried out against the following 20 drugs Cefuroxime (CXM), Ceftriaxone (CRO), Cefotaxime (CTX), Piperacillin (PIP), Sulphamethoxazole (SXT), Amoxicillin (AMC), Gentamicin (GN), Aztreonam (ATM), Ceftazidime (CAZ), Ciprofloxacin (CIP), Cefoxitin (FOX), Cefepime (FEP), Minocycline (MIN), Levofloxacin (LEV), Amikacin (AK), Piperacillin/Tazobactam (TZP), Imipenem (IPM), Meropenem (LEV), Amikacin (AK), Piperacillin/Tazobactam (TZP), Imipenem (IPM), Meropenem susceptibility is classified as either being Resistant (R), Intermediate (I), or Susceptible (S).

**S2 Table**. Antimicrobial resistance genes present in plasmids identified in *E. coli* plasmid predicted using the Resfinder database version 4.60.

## References

1. Centner TJ. Recent government regulations in the United States seek to ensure the effectiveness of antibiotics by limiting their agricultural use. Environment International. 2016. doi:10.1016/j.envint.2016.04.018

2. Prestinaci F, Pezzotti P, Pantosti A. Antimicrobial resistance: A global multifaceted phenomenon. Pathogens and Global Health. 2015. doi:10.1179/2047773215Y.0000000030

3. Murray CJ, Ikuta KS, Sharara F, Swetschinski L, Robles Aguilar G, Gray A, et al. Global burden of bacterial antimicrobial resistance in 2019: a systematic analysis. The Lancet. 2022;399. doi:10.1016/S0140-6736(21)02724-0

4. Gulumbe BH, Haruna UA, Almazan J, Ibrahim IH, Faggo AA, Bazata AY. Combating the menace of antimicrobial resistance in Africa: a review on stewardship, surveillance and diagnostic strategies. Biological Procedures Online. 2022. doi:10.1186/s12575-022-00182-y

5. Essack SY, Desta AT, Abotsi RE, Agoba EE. Antimicrobial resistance in the WHO African region: Current status and roadmap for action. Journal of Public Health (United Kingdom). 2017. doi:10.1093/pubmed/fdw015

6. World Health Organization. Global research agenda for antimicrobial resistance in human health. 2024. Available: https://books.google.com/books?hl=en&lr=&id=DFA4EQAAQBAJ&oi=fnd&pg=PR5&dq=WHO+bacterial+priority+pathogens+list,+2024:+Bacterial+pathogens+of+public+health+importance+to+guide+research,+development+and+strategies+to+prevent+and+control+antimicrobial+resistance&ots=SPKFEHatp0&sig=fkqw9UPGCAIvKuGzO6ljKm63Rz8

7. Kimani R, Wakaba P, Kamita M, Mbogo D, Mutai W, Ayodo C, et al. Detection of multidrug-resistant organisms of concern including Stenotrophomonas maltophilia and Burkholderia cepacia at a referral hospital in Kenya. journals.plos.orgR Kimani, P Wakaba, M Kamita, D Mbogo, W Mutai, C Ayodo, E Suliman, BN Kanoi, J GitakaPlos one, 2024•journals.plos.org. 2024;19. doi:10.1371/journal.pone.0298873

8. Rwigi D, Nyerere AK, Diakhate MM, Kariuki K, Tickell KD, Mutuma T, et al. Phenotypic and molecular characterization of β-lactamase-producing Klebsiella species among children discharged from hospital in Western Kenya. SpringerD Rwigi, AK Nyerere, MM Diakhate, K Kariuki, KD Tickell, T Mutuma, SN Tornberg, OO SogeBMC microbiology, 2024•Springer. 2024;24. doi:10.1186/s12866-024-03284-7

9. Omulo S, Oluka M, Achieng L, Osoro E, Kinuthia R, Guantai A, et al. Point-prevalence survey of antibiotic use at three public referral hospitals in Kenya. PLoS One. 2022;17. doi:10.1371/journal.pone.0270048

10. Bevan ER, Jones AM, Hawkey PM. Global epidemiology of CTX-M β-lactamases: Temporal and geographical shifts in genotype. Journal of Antimicrobial Chemotherapy. 2017;72. doi:10.1093/jac/dkx146

11. Mostafa HH. An evolution of Nanopore next-generation sequencing technology: implications for medical microbiology and public health. J Clin Microbiol. 2024;62. doi:10.1128/JCM.00246-24

12. Bogaerts B, Van den Bossche A, Verhaegen B, Delbrassinne L, Mattheus W, Nouws S, et al. Closing the gap: Oxford Nanopore Technologies R10 sequencing allows comparable results to Illumina sequencing for SNP-based outbreak investigation of bacterial pathogens. J Clin Microbiol. 2024;62. doi:10.1128/jcm.01576-23

13. Ye L, Liu X, Ni Y, Xu Y, Zheng Z, Chen K, et al. Comprehensive genomic and plasmid characterization of multidrug-resistant bacterial strains by R10.4.1 nanopore sequencing. Microbiol Res. 2024;283. doi:10.1016/j.micres.2024.127666

14. Sanderson ND, Hopkins KM V, Colpus M, Parker M, Lipworth S, Crook D, et al. Evaluation of the accuracy of bacterial genome reconstruction with Oxford Nanopore R10. 4.1 long-read-only sequencing. microbiologyresearch.orgND Sanderson, KMV Hopkins, M Colpus, M Parker, S Lipworth, D Crook, N StoesserMicrobial Genomics, 2024•microbiologyresearch.org. 2024;10. doi:10.1099/mgen.0.001246

15. Bonten M, Johnson JR, Van Den Biggelaar AHJ, Georgalis L, Geurtsen J, De Palacios PI, et al. Epidemiology of Escherichia coli Bacteremia: A Systematic Literature Review. Clinical Infectious Diseases. 2021. doi:10.1093/cid/ciaa210

16. Yeh TK, Lin HJ, Liu PY, Wang JH, Hsueh PR. Antibiotic resistance in Enterobacter hormaechei. International Journal of Antimicrobial Agents. 2022. doi:10.1016/j.ijantimicag.2022.106650

17. CLSI. M100 Performance Standards for Antimicrobial Susceptibility Testing A CLSI supplement for global application.

18. Wick RR, Judd LM, Holt KE. Performance of neural network basecalling tools for Oxford Nanopore sequencing. Genome Biol. 2019;20. doi:10.1186/s13059-019-1727-y

19. Freire B, Ladra S, Parama JR. Memory-Efficient Assembly Using Flye. IEEE/ACM Trans Comput Biol Bioinform. 2022;19: 3564–3577. doi:10.1109/TCBB.2021.3108843

20. Li H, Durbin R. Fast and accurate long-read alignment with Burrows-Wheeler transform. Bioinformatics. 2010;26: 589–595. doi:10.1093/bioinformatics/btp698

21. Vaser R, Sović I, Nagarajan N, Šikić M. Fast and accurate de novo genome assembly from long uncorrected reads. Genome Res. 2017;27: 737–746. doi:10.1101/gr.214270.116

22. Seemann T. Prokka: Rapid prokaryotic genome annotation. Bioinformatics. 2014;30. doi:10.1093/bioinformatics/btu153

23. Jolley KA, Bray JE, Maiden MCJ. Open-access bacterial population genomics: BIGSdb software, the PubMLST.org website and their applications. Wellcome Open Res. 2018;3. doi:10.12688/wellcomeopenres.14826.1

24. Jain C, Rodriguez-R LM, Phillippy AM, Konstantinidis KT, Aluru S. High throughput ANI analysis of 90K prokaryotic genomes reveals clear species boundaries. Nat Commun. 2018;9. doi:10.1038/s41467-018-07641-9

25. Beghain J, Bridier-Nahmias A, Nagard H Le, Denamur E, Clermont O. ClermonTyping: An easy-to-use and accurate in silico method for Escherichia genus strain phylotyping. Microb Genom. 2018;4. doi:10.1099/mgen.0.000192

26. Joensen KG, Tetzschner AMM, Iguchi A, Aarestrup FM, Scheutz F. Advances in Molecular Serotyping and Subtyping of Escherichia coli. frontiersin.org. 2015;53: 2402–2403. doi:10.1128/JCM.00008-15

27. Carattoli A, Zankari E, Garciá-Fernández A, Larsen MV, Lund O, Villa L, et al. PlasmidFinder and pMLST: in silico detection and typing of plasmid. Antimicrob Agents Chemother. 2014;58.

28. Robertson J, Nash JHE. MOB-suite: software tools for clustering, reconstruction and typing of plasmids from draft assemblies. Microb Genom. 2018;4. doi:10.1099/mgen.0.000206

29. Maxime B, Sébastien L. GenoFig: a user-friendly application for the visualisation and comparison of genomic regions. Bioinformatics. 2024 [cited 2 Jan 2025]. Available: https://academic.oup.com/bioinformatics/advance-article-abstract/doi/10.1093/bioinformatics/btae372/7693070

30. Grant J, Enns E, Marinier E, Mandal A, Herman KE, Chen C, et al. Proksee: in-depth characterization and visualization of bacterial genomes. Nucleic Acids Res. 2023 [cited 2 Jan 2025]. Available: https://academic.oup.com/nar/article-abstract/51/W1/W484/7151341

31. Mezzatesta ML, Gona F, Stefani S. Enterobacter cloacae complex: Clinical impact and emerging antibiotic resistance. Future Microbiology. 2012. doi:10.2217/fmb.12.61

32. Weldetensae MK, Weledegebriel MG, Nigusse AT, Berhe E, Gebrearegay H. Catheter-Related Blood Stream Infections and Associated Factors Among Hemodialysis Patients in a Tertiary Care Hospital. Infect Drug Resist. 2023;16. doi:10.2147/IDR.S409400

33. Mathers AJ, Peirano G, Pitout JDD. The role of epidemic resistance plasmids and international high-risk clones in the spread of multidrug-resistant Enterobacteriaceae. Clin Microbiol Rev. 2015;28. doi:10.1128/CMR.00116-14

34. Johnson TJ, Elnekave E, Miller EA, Munoz-Aguayo J, Figueroa CF, Johnston B, et al. Phylogenomic Analysis of Extraintestinal Pathogenic Escherichia coli Sequence Type 1193, an Emerging Multidrug-Resistant Clonal Group. Antimicrob Agents Chemother. 2019;63. doi:10.1128/AAC.01913-18

35. Soliman AM, Maruyama F, Zarad HO, Ota A, Nariya H, Shimamoto T, et al. Emergence of a multidrug-resistant enterobacter hormaechei clinical isolate from egypt co-harboring mcr-9 and blavim-4. Microorganisms. 2020;8. doi:10.3390/microorganisms8040595

36. Kananizadeh P, Oshiro S, Watanabe S, Iwata S, Kuwahara-Arai K, Shimojima M, et al. Emergence of carbapenem-resistant and colistin-susceptible Enterobacter cloacae complex co-harboring bla IMP-1 and mcr-9 in Japan. BMC Infect Dis. 2020;20. doi:10.1186/s12879-020-05021-7

37. Dimitriu T. Evolution of horizontal transmission in antimicrobial resistance plasmids. Microbiology (United Kingdom). 2022. doi:10.1099/mic.0.001214

38. Berglund B. Environmental dissemination of antibiotic resistance genes and correlation to anthropogenic contamination with antibiotics. Infect Ecol Epidemiol. 2015;5. doi:10.3402/iee.v5.28564

39. Murray L, Hayes A. Co-selection for antibiotic resistance by environmental contaminants. Nature. 2024 [cited 31 Dec 2024]. Available: https://www.nature.com/articles/s44259-024-00026-7

40. Nauclér P, Huttner A, van Werkhoven CH, Singer M, Tattevin P, Einav S, et al. Impact of time to antibiotic therapy on clinical outcome in patients with bacterial infections in the emergency department: implications for antimicrobial stewardship. Clinical Microbiology and Infection. 2021. doi:10.1016/j.cmi.2020.02.032

41. Maharjan S, Gallagher P, Gautam M, Joh HS, Sujan MJ, Aboushady AT, et al. Recording and Reporting of Antimicrobial Resistance (AMR) Priority Variables and Its Implication on Expanding Surveillance Sites in Nepal: A CAPTURA Experience. Clinical Infectious Diseases. 2023;77. doi:10.1093/cid/ciad581

42. Porter G, Kotwani A, Bhullar L, Joshi J. Over-the-counter sales of antibiotics for human use in India: The challenges and opportunities for regulation. Med Law Int. 2021;21. doi:10.1177/09685332211020786

43. Akpan MR, Isemin NU, Udoh AE, Ashiru-Oredope D. Implementation of antimicrobial stewardship programmes in African countries: a systematic literature review. Journal of Global Antimicrobial Resistance. 2020. doi:10.1016/j.jgar.2020.03.009

