## Supplemental Table 1 for "Genomic characterization of *Escherichia coli* and *Enterobacter hormaechei* clinical isolates from a tertiary healthcare facility in Kenya"

| **Sample** | **Species** | **CXM**  **(30µg)** | **CRO**  **(30µg)** | **CTX**  **(30µg)** | **PRL**  **(30µg)** | **SXT**  **(25µg)** | **AMC**  **(10µg)** | **CN**  **(10µg)** | **ATM**  **(30µg)** | **CAZ**  **(30µg)** | **CIP**  **(5µg)** | **FOX**  **(30µg)** | **FEP**  **(30µg)** | **MH**  **(30µg)** | **LEV**  **(5µg)** | **AK**  **(30µg)** | **TZP**  **(110µg)** | **IPM**  **(10µg)** | **MEM**  **(10µg)** | **TET**  **(30µg)** | **NA**  **(10µg)** | **AMP**  **(10µg)** |
| --- | --- | --- | --- | --- | --- | --- | --- | --- | --- | --- | --- | --- | --- | --- | --- | --- | --- | --- | --- | --- | --- | --- |
| S03 | *E. hormaechei* | R | R | R | R | R | R | R | R | R | R | R | R | I | R | S | R | S | S | R | R | R |
| S08 | *E. hormaechei* | R | R | R | R | R | R | R | R | R | I | R | R | I | I | I | R | R | R | S | R | R |
| S09 | *E. hormaechei* | R | R | R | R | R | R | R | R | R | R | R | R | R | R | I | I | S | S | R | R | R |
| S24 | *E.coli* | R | R | R | R | S | S | R | R | R | S | R | I | S | S | I | R | R | R | S | I | R |
| S26 | *E.coli* | R | R | R | R | R | I | R | R | S | R | S | R | R | R | S | S | I | S | R | R | R |
| S27 | *E. coli* | R | R | R | R | R | R | S | S | R | R | S | R | R | R | I | R | R | S | R | R | R |
| S29 | *E. coli* | R | R | R | R | S | R | S | R | R | R | I | R | I | R | S | S | S | S | R | R | R |
