## Supplemental Table 2 for "Genomic characterization of *Escherichia coli* and *Enterobacter hormaechei* clinical isolates from a tertiary healthcare facility in Kenya"

| Sample ID | Plasmid ID | Plasmid replicon | Length (bp) | Acquired AMR genes |
| --- | --- | --- | --- | --- |
| S29 | pS29_003 | RepA | 287,498 | *aadA1, dfrA1,blaTEM-148, dfrA1, blaTEM-1B, sul1, qacE* |
|  | pS29_002 | lncR | 55,769 | *qacE, sul1, mph(A), ARR-3, blaDHA-1, qnrB4, blaOXA-1, qacE, blaDHA-27, aac(6')-Ib-cr, catB3* |
